# Identification of the Clostridial cellulose synthase and characterization of the cognate glycosyl hydrolase, CcsZ

**DOI:** 10.1101/837344

**Authors:** William Scott, Brian Lowrance, Alexander C. Anderson, Joel T. Weadge

**Affiliations:** Department of Biology, Wilfrid Laurier University 75 University Ave W. Waterloo, ON N2L 3C5

**Author notes:** Corresponding author: Joel T. Weadge, Department of Biology, Wilfrid Laurier University, 75 University Ave W. Waterloo, ON N2L3C5. These authors contributed equally to this work (William Scott, Brian Lowrance, Alexander C. Anderson). Department of Molecular and Cellular Biology, University of Guelph, 50 Stone Rd. E Guelph, ON N1G 2W1.

## Abstract

Biofilms are community structures of bacteria enmeshed in a self-produced matrix of exopolysaccharides. The biofilm matrix serves numerous roles, including resilience and persistence, making biofilms a subject of research interest among persistent clinical pathogens of global health importance. Our current understanding of the underlying biochemical pathways responsible for biosynthesis of these exopolysaccharides is largely limited to Gram-negative bacteria. Clostridia are a class of Gram-positive, anaerobic and spore-forming bacteria, and include the important human pathogens *Clostridium perfringens, Clostridium botulinum, and Clostridioides difficile*, among numerous others. Clostridia have been reported to form biofilms composed of cellulose, although the specific loci which encode the cellulose synthase have not been identified. Here, we report the discovery of a gene cluster, which we named *ccsABZCD*, among selected bacteria within class Clostridia that appears to encode a synthase complex responsible for polymerization, modification, and export of an O-acetylated cellulose exopolysaccharide. To test this hypothesis, we subcloned the putative glycoside hydrolase CcsZ and solved the X-ray crystal structure of both apo- and product-bound CcsZ. Our results demonstrate that CcsZ is in fact an *endo*-acting cellulase belonging to glycoside hydrolase family 5 (GH-5). This is in contrast to the Gram-negative cellulose synthase, which instead encodes BcsZ, a structurally distinct GH-8. We further show CcsZ is capable of hydrolysis of the soluble mock substrate carboxymethylcellulose (CMC) with a pH optimum of 4.5. The data we present here serves as an entry point to an understanding of biofilm formation among class Clostridia and allowed us to predict a model for Clostridial cellulose synthesis.

**Author summary:** Biofilms are communities of microorganisms that enmesh themselves in a protective matrix of elf-produced polysaccharide materials. Biofilms have demonstrated roles in both virulence and persistence among bacterial pathogens of global health importance. The class Clostridia are a polyphyletic grouping of primarily Gram-positive, anaerobic and spore-forming bacteria which contain the important and well-studied human pathogens *Clostridioides difficile, Clostridium botulinum,* and *Clostridium perfringens,* among others. Bacteria belonging to class Clostridia have been anecdotally reported to form biofilms made of cellulose, although the molecular mechanisms governing their production has not before been described. In this work, we identify a gene cluster, which we name *ccsABZHI*, for the <u>C</u>lostridial <u>c</u>ellulose <u>s</u>ynthase, which bears remarkable similarity to molecular machinery required for the production of cellulose biofilms in other Gram-negative bacteria. We further biochemically characterize one of these enzymes, CcsZ, a predicted endoglucanase which we predicted from our model should cleave cellulose exopolysaccharides. We show that CcsZ is in fact capable of this activity and belongs to a broader family of glycoside hydrolases with unexpected taxonomic diversity. Our work represents an entry point towards an understanding of the molecular mechanisms governing cellulose biofilm formation in Gram-positive bacteria.

## Introduction

Biofilms are communities of microorganisms that reside in an extracellular matrix. This extracellular matrix is produced by the community itself and is composed largely of secreted exopolysaccharides. Biofilms are among the most successful and widely distributed forms of life on Earth, enabling bacteria to adhere to, colonize, and persist on a wide variety of surfaces or interfaces. The production of the biofilm extracellular matrix represents a significant resource cost to the organisms producing it, rationalized by the many benefits the biofilm matrix confers. For example, the biofilm matrix serves in resource capture [1, 2], accelerated cell growth [3], and the tolerance of stressors or disinfectants, such as shearing forces [2, 3], desiccation [2, 4], antimicrobial compounds [2, 5], extreme temperature [2, 5], or sanitizing agents [2,5,6]. Production of an extracellular matrix composed of the exopolysaccharide cellulose has been described in many species of clinical or economic importance, including *Acetobacter* [7]*, Clostridium* (previously *Sarcina*) [7, 8]*, Rhizobium* [7]*, Agrobacterium* [7]*, Cronobacter* [9]*, Salmonella* [10], and *Pseudomonas* [11], among others. The production of a linear β-(1-4)-glucan polymer by these bacteria is accompanied by the derivatization of the polysaccharide into novel biomaterials suited to the needs of the particular organism producing it and the niche being colonized [12]. These exopolysaccharide modifications enhance bacterial persistence and have been identified as a virulence factor under some circumstances [13].

At present, an understanding of bacterial cellulose biosynthesis has been largely limited to the study of Gram-negative model organisms. The biosynthesis and export of exopolysaccharides in bacteria is carried out by synthase-dependent systems, encoded on partially conserved operons [14]. Pertaining to cellulose, the Gram-negative bacterial cellulose synthesis (Bcs) complex is encoded by the *bcsABCZ* locus, and represents the essential biosynthetic machinery required for production of the β-(1-4)-glucan polymer [15] that is subsequently derivatized by at least three distinct accessory cellulose modification systems in Gram-negative bacteria [11,16,17]. The proposed functions of the *bcsABCZ* gene products were largely inferred from structural and functional studies of other Gram-negative exopolysaccharide systems, *e.g.* the poly-β-(1-6)-N-acetyl-D-gluccoasmine (PNAG) biosynthetic system from *Escherichia coli* [18] and *Staphylococcal* species [19, 20] or the alginate biosynthesis pathway from *Pseudomonas aeruginosa* [21]. Subsequently, structural and functional studies of the Bcs gene products have since provided insight to their functional roles. In these systems, biosynthesis of the glucan polymer is carried out by the family 2 glycosyltransferase (GT-2) BcsA [22]. The processive action of BcsA uses UDP-glucose as a substrate to successively add the sugars to the growing chain, while concomitantly transporting the elongated chain through a pore in BcsA to the periplasm, where it interacts with the carbohydrate-binding module (CBM) contained within BcsB [22, 23]. Although the mechanism whereby the nascent cellulose chain is exported from the cell has not been directly elucidated, the BcsC protein is proposed to contain both a β-barrel domain and a tetratricopeptide repeat (TPR) domain to facilitate both the assembly of the Bcs complex and the efficient export of the polymer to the extracellular space [14]. This model is in line with the alginate biosynthesis pathway, where AlgE forms a β-barrel necessary for exopolysaccharide export, and AlgK is a TPR-containing protein [24, 25]. In the alginate system, release of the polymer from the cell is accomplished by the action of an alginate lyase AlgL [26]. In the Bcs system, this role is accomplished by the family 8 glycoside hydrolase (GH-8) BcsZ [27, 28].

Exopolysaccharide biosynthesis in Gram-positive bacteria, although still poorly understood, has been best studied in the model organisms *Staphylococcus aureus* and *Staphylococcus epidermidis*, which produce a biofilm composed of PNAG produced by the *icaADBC* locus [29]. Structural and functional studies of the *icaADCB* gene products have suggested that PNAG biosynthesis in *S. aureus* and *S. epidermidis* occurs through an analogous, but distinct, synthase-dependent mechanism to the Gram-negative PNAG synthase, encoded by the *pgaABCD* locus [29]. For example, the lack of both a TPR-containing region and β-barrel in the Ica complex is rationalized by the absence of an outer membrane, whereby translocation across the cytoplasmic membrane of Gram-positive bacteria is sufficient for extracellular release of the PNAG polysaccharide [29]. Additionally, enzymatic modification of the PNAG polymer by the PgaB deacetylase occurs in the periplasm of Gram-negative bacteria [30]; the IcaB de-N-acetylase is instead an extracellular protein [31].

Evidence for the production of a cellulose biofilm by the Gram-positive and strictly anaerobic bacterium *Clostridium ventriculi* (formerly *Sarcina ventriculi*) was first reported by Canale-Parole *et al.* (1961, Ref. 8). To date, numerous Clostridia have been reported to form biofilms, including the important human pathogens *Clostridioides difficile* (formerly *Clostridium difficile*), *Clostridium perfringens* and *Clostridium botulinum*, a phenomenon that has been reviewed by others [32]. Only just recently was the Pel polysaccharide biosynthetic gene cluster identified in Gram-positive bacteria, including *Clostridia* [33]. However, the loci encoding other likely exopolysaccharide synthase components have not been identified in any of these species, and the specific requirements for biofilm formation in Clostridia remain largely unexplored.

To identify the Clostridial cellulose synthase, we surveyed available sequence databases for predicted orthologues of the Gram-negative cellulose synthase components. We identified a 5-gene locus, apparently conserved within class Clostridia, which we designate herein as the <u>C</u>lostridial <u>c</u>ellulose <u>s</u>ynthase (*ccsABZHI*) to distinguish it from the Gram-negative Bcs locus, based on sequence annotation and homology modelling. Based upon this apparent homology, the locus appears to encode a homologous glycosyltransferase (CcsA) and glycoside hydrolase (CcsZ), and a protein of unknown function (CcsB), with the notable absence of an equivalent to the Gram-negative outer membrane component (BcsC). Additionally, sequence homology and modelling predicted that *ccsH* and *ccsI* may encode a complex for O-acetylation of the cellulose exopolysaccharide. To investigate our hypothesis that *ccsABZHI* is responsible for cellulose synthesis, we selected the putative glycoside hydrolase (CcsZ) for cloning and expression, as this protein would be expected to be soluble and for which a structural and functional characterization would be straightforward. Accordingly, we heterologously expressed CcsZ in an *E. coli* host as a His-tagged fusion protein and purified it to apparent homogeneity. Structure determination revealed CcsZ is a family 5 glycoside hydrolase with an (α/β)_8_ TIM barrel fold, in contrast to available structures of the Gram-negative cellulose synthase component BcsZ from *E. coli* and *P. putida*, both GH-8 enzymes with an (α/α)_6_ barrel fold. Structural comparison and analysis showed CcsZ has an unusual architecture for substrate accommodation among GH-5, which was further supported by its product-bound structure. CcsZ demonstrated activity on the mock substrate carboxymethylcellulose (CMC), with a pH optimum of 4.5, typical of GH-5 cellulases, and was either not active or only minimally active on other common GH-5 substrates. Furthermore, CcsZ demonstrated maximal activity on β-glucan, a soluble cellulose-analog substrate. The model of the Clostridial cellulose synthesis system we propose here represents an entry point to an understanding of cellulose biofilm formation by Gram-positive bacteria and enables future structural and functional studies of other Clostridial cellulose synthesis subunits.

## Materials and Methods

### Identification of *ccsABZHI*

Initial identification of the Clostridial cellulose synthase was carried out by searching the NCBI database (https://www.ncbi.nlm.nih.gov) for sequences annotated as BcsA orthologues or by BlastP search (https://blast.ncbi.nlm.nih.gov/Blast.cgi) using the *E. coli* K12 BcsA sequence as a query, and filtering results for sequences from Clostridia only. We then identified other putative *ccs* ORFs based upon their proximity to BcsA orthologues and through functional annotation, BlastP searches, and homology modelling on the PHYRE2 server [34] (http://www.sbg.bio.ic.ac.uk/phyre2/) ascertained their likely biochemical roles. Sequence alignments and identities were calculated using the BlastP tool. The topology of CcsB was predicted using the TMHHM server, v.2.0 (32; http://www.cbs.dtu.dk/services/TMHMM/).

### Cloning

The full-length sequence of the hypothetical exopolysaccharide synthesis-associated glycoside hydrolase from *C. difficile* (GenPept accession WP_077724661.1), which we designated *ccsZ*, was predicted to have an N-terminal signal sequence according to the PSort server [36] that remains as a single transmembrane anchor in the mature protein. A *ccsZ* plasmid construct lacking the transmembrane region (amino acids 1-29) was synthesized by BioBasic, Inc. (Markham, ON) in a pET21 vector background so that the expressed soluble portion of the protein (amino acids 30-340) would contain a C-terminal hexahistidine tag (pCcsZ-His_6_).

### CcsZ expression and purification

The expression host, *E. coli* BL21-CodonPlus, was transformed with pCcsZ-His_6_ and cells were grown in rich media (32 g tryptone, 20 g yeast extract and 10 g NaCl) containing ampicillin (100 μg mL^-1^) and chloramphenicol (34 μg mL^-1^) at 37° C with shaking. Expression was induced with IPTG (1 mM) after cultures reached an OD_600_ of at least 0.6 and growth was allowed to continue 16 – 20 h at 23°C. The cells were collected by centrifugation at 5000 x *g* for 15 min and 4°C. The pellet was resuspended in buffer A (50 mM Tris-HCl pH 7.5, 300 mM NaCl) and cells were lysed with a cell disruptor (Constant Systems) operating at 17 kpsi. Cell lysate was separated by centrifugation at 28000 x *g* for 45 min and 4°C and the supernatant was loaded onto a column containing 2 mL of Ni-NTA agarose (Qiagen) equilibrated with buffer A for 1 h for purification by nickel affinity chromatography. The Ni-NTA column was washed in 50 mL each of buffer A, and buffer A with the addition of 20 mM imidazole-HCl, followed by a wash in 50 mL of buffer A with the addition of 50 mM imidazole-HCl. The protein was eluted from nickel affinity chromatography columns using 25 mL of buffer A with the addition of 500 mM imidazole-HCl. Secondary purification was performed by anion exchange chromatography using a HiTrap Q FF 5 mL prepacked column (GE Healthcare) installed on an Akta Start instrument (GE Healthcare). Protein eluted from Ni-NTA resin was dialyzed against anion buffer A (50 mM sodium phosphate pH 7.5) for 12 – 16 h and passed over the column three times. CcsZ-His_6_ was eluted from the column using a gradient of 0-100 % anion buffer B (50 mM sodium phosphate pH 7.5, 1 M NaCl). The purity of protein obtained from both nickel affinity chromatography and anion exchange chromatography was routinely assessed using SDS-PAGE. Where needed, purified protein was concentrated in a centrifugal filter unit (Pall Corporation) with a nominal MWCO of 10,000 Da.

### Crystallization and structure determination

Crystallization of CcsZ was carried out at 20°C using the hanging drop vapour diffusion method. Apo-CcsZ crystals were obtained from drops containing 100 mM Tris-HCl pH 8.5, 200 mM calcium acetate, and 25% (v/v) poly(ethylene glycol) 4000 in a 1:1 ratio with CcsZ protein concentrated to 12.5 mg/mL. Product-bound crystals were obtained from drops containing 100 mM MES pH 6.0, 200 mM calcium acetate, and 20% (v/v) poly(ethylene glycol) 8000 (PEG 8000) in a 1:1 ratio with CcsZ concentrated to 12.5 mg/mL. Cellotriose was introduced to the drop at a concentration of 5 mM following crystal growth and to cryoprotectants (crystallization buffer supplemented to 30% (v/v) PEG 8000) just prior to harvesting and vitrification in liquid nitrogen. Data were collected on beamline 08-ID1 at the Canadian Light Source synchrotron. A total of 1250 images of 0.2° Δφ oscillations were collected using incident radiation with a wavelength of 1.0 Å for the apo-CcsZ crystal and a total of 1800 images of 0.2° Δφ oscillations were collected using incident radiation with a wavelength of 1.0 Å for the product-bound crystal. Collected data was processed with XDS [37]. A *P* 2_1_ space group was determined with a single copy of CcsZ in the asymmetric unit using POINTLESS, then scaled using SCALA and data reduction performed using CTRUNCATE [38]. The processed data was solved using the molecular replacement technique with the Phaser tool in PHENIX [39] using *Tm*Cel5A (PDB ID 3AMD) as a search model. Structural refinement was performed using iterative rounds of automated refinement using the Refine tool in PHENIX, followed by manual refinement and checking of waters using Coot [40]. Refinement progress was monitored by the reduction and convergence of *R*_work_ and *R*_free_. Ligand fitting was performed by LigandFit in PHENIX, followed by calculation of an omit map by Polder in PHENIX. Ligand real-space refinement was performed in Coot using the Polder map, and validation was performed using MolProbity in PHENIX. Structure interface analysis was performed using the PDBePISA server (https://www.ebi.ac.uk/pdbe/pisa/pistart.html) [41]. Figures were prepared with PyMOL.

### Glycoside hydrolase activity assays

Activity was initially assessed qualitatively by spotting proteins on an agar overlay containing dissolved carbohydrates. Agar plates were prepared by heating a solution of 1.5% (w/v) agar and 0.8% (w/v) carbohydrate to 100°C for 10 min. Carbohydrates tested with the agar overlay include CMC, xylan, and hydroxyethylcellulose (Sigma). Spotted on the agar plates were 0.5 mg each of purified CcsZ, cellulase cocktail from *Trichoderma reesei* (Sigma; C8546), and bovine serum albumin (BSA; BioShop). BSA served as the negative control and the *T. reesei* cellulase cocktail served as the positive control. The plates were incubated either 1 h or 24 h, as specified, at 37°C. Following incubation, plates were stained in 5 mL of a 0.1% (w/v) solution of Congo Red for 1 h and destained in three 5 mL volumes of a 1 M solution of NaCl for 1 h total.

The activity of CcsZ was quantitatively monitored using the dinitrosalicylic acid (DNS) method. Substrates tested included carboxymethylcellulose (CMC), xylan, arabinoxylan, β-glucan, lichenin and xylan. All carbohydrates not purchased from Sigma as noted above were supplied by Megazyme. All substrates were tested at 8.3 g/L. CcsZ (20 μM) was mixed with substrate in assay buffer (50 mM sodium acetate pH 4.5) for 30 min at 37°C. Reactions were stopped with the addition of 56.25 μL of DNS reagent (1% w/v dinitrosalicylic acid, 4 mM sodium sulfite, 250 mM NaOH) and were heated at 90°C for 15 min. A volume of 18.75 μL of (40% w/v) sodium potassium tartrate was added to stabilize the DNS reaction. The resulting solutions were measured for absorbance at 575nm in a Cytation5 imaging microplate reader (BioTek). Absorbance values were related to millimolar reducing sugar equivalents using a standard curve of D-glucose (BioShop) measured using the same DNS method. Buffers used for the pH assay included sodium acetate (pH 4.0-5.5), MES (pH 5.5-7), sodium phosphate (pH 7.0-8.0), Tris-HCl (pH 8.0-9.0), and CHES (pH 9.0-10) in increments of 0.5 pH units with each buffer at 50 mM final concentration.

### LC-MS

Liquid chromatography–mass spectrometry analyses were performed at the Mass Spectrometry Facility of the Advanced Analysis Centre, University of Guelph on an Agilent 1200 HPLC liquid chromatograph interfaced with an Agilent UHD 6530 Q-TOF mass spectrometer. A C18 column (Agilent Extend-C18 50 mm x 2.1 mm 1.8 µm) was used for chromatographic separation. The following solvents were used for separation: water with 0.1% (v/v) formic acid (A) and acetonitrile with 0.1% (v/v) formic acid (B). The initial mobile phase conditions were 10% B, hold for 1 min, and then increase to 100% B in 29 min. These steps were followed by a column wash at 100% B for 5 min and a 20 min re-equilibration. The mass spectrometer electrospray capillary voltage was maintained at 4.0 kV and the drying gas temperature at 250° C with a flow rate of 8 L/min. Nebulizer pressure was 30 psi and the fragmentor was set to 160. Nozzle, skimmer and octapole RF voltages were set at 1000 V, 65 V and 750 V, respectively. Nitrogen (>99%) was used as the nebulizing, drying and collision gas. The mass-to-charge ratio was scanned across the m/z range of 50-3000 m/z in 4 GHz (extended dynamic range) positive and negative ion modes. Data were collected by data independent MS/MS acquisition with an MS and MS/MS scan rate of 1.41 spectra/sec. The acquisition rate was set at 2 spectra/s. The mass axis was calibrated using the Agilent tuning mix HP0321 (Agilent Technologies) prepared in acetonitrile. Mass spectrometer control, data acquisition and data analysis were performed with MassHunter® Workstation software (B.04.00).

## Results and Discussion

### Identification of the Clostridial cellulose synthase

We searched the NCBI Protein database for hypothetical proteins that were functionally annotated as cellulose synthases, or which showed similarity to known Gram-negative BcsA proteins among the Clostridia. We identified sequences functionally annotated as putative cellulose synthase catalytic subunits in *C. difficile* (GenBank GAX65516.1)*, Clostridium vincentii* (PRR81681.1), *Clostridium chromireducens* (OPJ65957.1), and *Clostridium oryzae* (OPJ56113.1), among others (Fig. 1; red). The functional annotations are based on sequence similarity to the GT-2 domain of *E. coli* BcsA (UniProt P37653) and the presence of a predicted PilZ domain for binding c-di-GMP, a hallmark of synthase-dependent exopolysaccharide catalytic subunits. These features were a promising indication these proteins were likely true orthologues of BcsA. Homology modelling of these hypothetical proteins also suggested they belong to GT-2, with PHYRE2 selecting the structure of BcsA from *Rhodobacter sphaeroides* (PDB ID 4HG6) as the top scoring homologue and the template for homology modelling in all cases.

**Figure 1.**
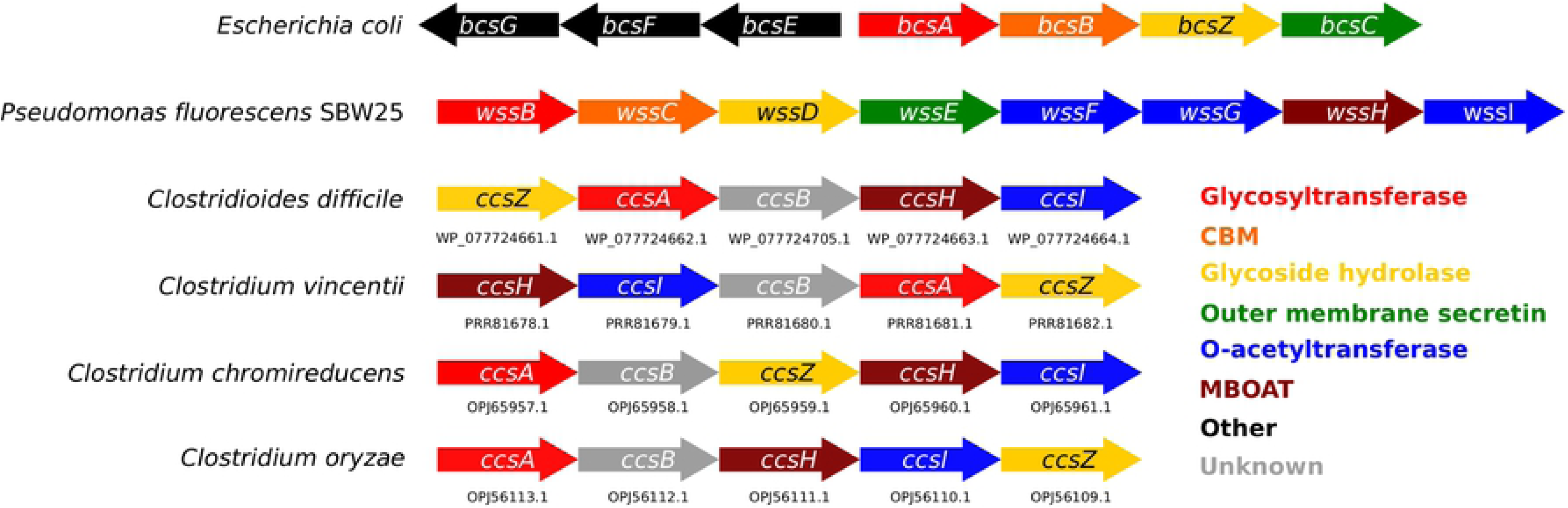
The conserved *ccsABZHI* locus. A set of 5 genes proposed to be involved in O-acetylated cellulose exopolysaccharide biosynthesis was identified in *Clostridioides difficile* and appears conserved in other bacteria of class Clostridia. These genes encode apparent orthologues of the Gram-negative cellulose synthase complex, including equivalents to the glycosyltransferase BcsA and the glycosyl hydrolase BcsZ, in addition to a putative MBOAT and O-acetyltransferase that may O-acetylate cellulose exopolysaccharides. The GenBank accession codes for *ccs* genes we identified are listed below each gene.

We subsequently examined the loci adjacent to CcsA in each organism listed above to identify if other cellulose synthase subunits were present in a similar orientation to the *bcsABZC* operon found in Gram-negative bacteria (Fig. 1). Although the orientation of specific genes varied between the genomes we surveyed, in all cases we also identified a putative glycoside hydrolase, annotated as an *endo*-glucanase precursor protein, which we presumed to be the functional equivalent of BcsZ and which we denoted CcsZ (Fig 1; yellow). In addition, we identified a highly conserved protein of unknown function, which we denote herein as CcsB (Fig. 1; gray). Although it is tempting to speculate that these CcsB proteins may serve the functional equivalent to BcsB, a BlastP search of these sequences returned only other proteins of unknown function found in members of class Clostridia and provided no evidence of homology to BcsB. Homology modelling predicted these CcsB sequences to be related to exo-β-agarases, although we interpreted this with skepticism due to very limited coverage and identity to the CcsB sequence against the agarase domain fold (*i.e.*, < 40% coverage with < 20% identity in all cases). To understand the localization of CcsB, we analyzed the sequence with the TMHMM bioinformatics tool that predicted the sequence contains two transmembrane helices of 19 and 22 residues in length, with these helices very near to each terminus and connected by an approximately 300 amino acid extracellular domain. As expected, no protein was identified with predicted homology to BcsC, or that contained a predicted TPR or β-barrel domain that would be required for export across an outer membrane; a function mandated only in Gram-negative bacteria (Fig. 1; green).

We also located two conserved loci that were always found adjacent to *ccsABZ*, which we named *ccsH* and *ccsI* (Fig. 1; blue and brown). CcsH was often annotated as a membrane-bound O-acetyltransferase (MBOAT) protein, further supported by homology modelling. Although CcsI was annotated as a protein of unknown function in all cases, BlastP searches and homology modelling using CcsI as a query suggested that *ccsI* encodes a putative O-acetyltransferase protein from the GDSL family. Taken together, this finding indicated to us that *ccsABZHI* probably encodes a cellulose synthase that produces an O-acetylated cellulose exopolysaccharide.

O-acetylation of cellulose exopolysaccharides are one enzymatic modification identified in Gram-negative bacteria, first described by Spiers *et al*., where biosynthesis in the model organism *P. fluorescens* SBW25 is proposed to be carried out by the *wssABCDEFGHIJ* operon [11]. In this system, *wssBCDE* are proposed functional equivalents to *bcsABZC*, with *wssAJ* predicted to serve in cellular localization of the synthase complex, and *wssFGHI* predicted to serve in cellulose O-acetylation [11] in an analogous fashion to the alginate acetyltransferases *algXFIJ* [42]. The *ccsH* and *ccsI* genes are annotated as an MBOAT family protein and an AlgX_N_like/AlgJ O-acetyltransferase, respectively, in Clostridia; thereby suggesting they may be homologous to known Gram-negative exopolysaccharide O-acetyltransferases. Our own homology searches (BLAST Global Align and PHYRE2 modelling) confirm these predictions, as CcsH and CcsI modelled with *≥* 99% confidence and ≥ 74% sequence coverage against the structure of the MBOAT protein, DltB, and the O-acetyltransferase protein, PatB1, from *B. cereus*, respectively.

The role of such MBOAT and O-acetyltransferase proteins in similar pathways for the O-acetylation of secondary cell wall polysaccharides (SCWPs) has been demonstrated previously [43]. In this system, the MBOAT protein PatA1 is probably responsible for translocation of an acetyl group from a cytoplasmic donor, likely acetyl-coenzyme A (CoA-Ac) to the extracellular face of the cytoplasmic membrane [44]. PatB1, a peripheral membrane protein, then catalyzes the transfer of the acetyl group to terminal SCWP GlcNAc residues, a modification enabling proper assembly of the S-layer [43]. This system also closely resembles the well-characterized peptidoglycan O-acetyltransferase systems encoded by OatA in *Streptococcus pneumoniae* and PatA/PatB in various Gram-negative bacteria, which has been reviewed extensively by Scyhanatha *et al* [44]. It is also worth noting that a similar phenomenon is observed in the Gram-positive PNAG synthase, where O-succinylation of PNAG has been proposed to be carried out by IcaC, an apparent member of the acyltransferase family 3 (AT-3), most closely resembling the peptidoglycan O-acetyltransferase OatA from *S. aureus* [29]. Taken in this context, our results further establish a link between the pathways responsible for O-acetylation of cell wall-associated polysaccharides and secreted exopolysaccharides.

Based on analogy to existing models for exopolysaccharide synthesis and modification in both Gram-negative and Gram-positive bacteria, our analysis of *ccsABZHI* enabled us to predict a model for the O-acetylated cellulose synthase in selected Clostridia (Fig. 2C). This proposed cellulose synthase shares important structural characteristics with both the Gram-negative cellulose synthase and the Gram-positive PNAG synthase (Fig. 2). Our model of the Clostridial cellulose synthase resembles the Gram-negative cellulose synthase in that the catalytic subunit (CcsA) possesses the hallmark GT-2 domain which would be responsible for assembling the linear β-(1,4) glucan polymer, in addition to a PilZ domain responsible for c-di-GMP binding and regulatory control of synthase activity (Fig. 2, A and C). However, the remaining features of the Clostridial cellulose synthase appear structurally similar to the Gram-positive PNAG synthase encoded by the *icaABCD* locus. IcaA and IcaD are membrane proteins that together are responsible for the production of the PNAG polymer [45]. Interestingly, IcaA contains a canonical GT-2 domain, while IcaD is a smaller protein of unknown function, predicted to be a membrane protein with two TM helices, yet its expression was still necessary for maximal IcaA activity and correct PNAG synthesis [45]. Although CcsB and IcaD share a poor sequence alignment (> 16%), low sequence identity (31%), and no predicted homology, both of these proteins appear conserved in their respective gene clusters and contain two predicted terminal TM helices linked by a single extracellular domain, although CcsB is much larger than IcaD (*i.e.* 358 residues versus 101)[29, 45]. Thus, our model would predict that *ccsAB* is necessary and sufficient for cellulose polymerization at the cytoplasmic membrane as reported for *icaAD*. Furthermore, the glycoside hydrolase found on virtually all known exopolysaccharide synthesis gene clusters is present here as CcsZ. Additionally, our model accounts for the lack of an outer membrane component required for export only in Gram-negative bacteria, but does share the feature of a polysaccharide modification system (CcsH/CcsI) with homology to other extracellular carbohydrate acetyltransferase systems (WssH/WssI in *P. fluorescens* and AlgIJ in *Pseudomonas aeruginosa*), as well as the peptidoglycan O-acetyltransferases (PatA/PatB and the AT-3 containing OatA-like protein IcaC in *Staphylococcus* spp.) (Fig. 2B). The observation that CcsH/CcsI in the Clostridia is related to alginate and peptidoglycan O-acetyltransferases suggests an emerging trend of bacterial persistence for these O-acetyltransferase proteins, as O-acetylation of peptidoglycan confers resistance to lysozyme and is a virulence factor in Gram-positive bacteria [44], while O-acetylation of alginate contributes to biofilm architecture and adhesion, as well as improved bacterial resistance to antibiotics and opsonic phagocytosis [46–48].

**Figure 2.**
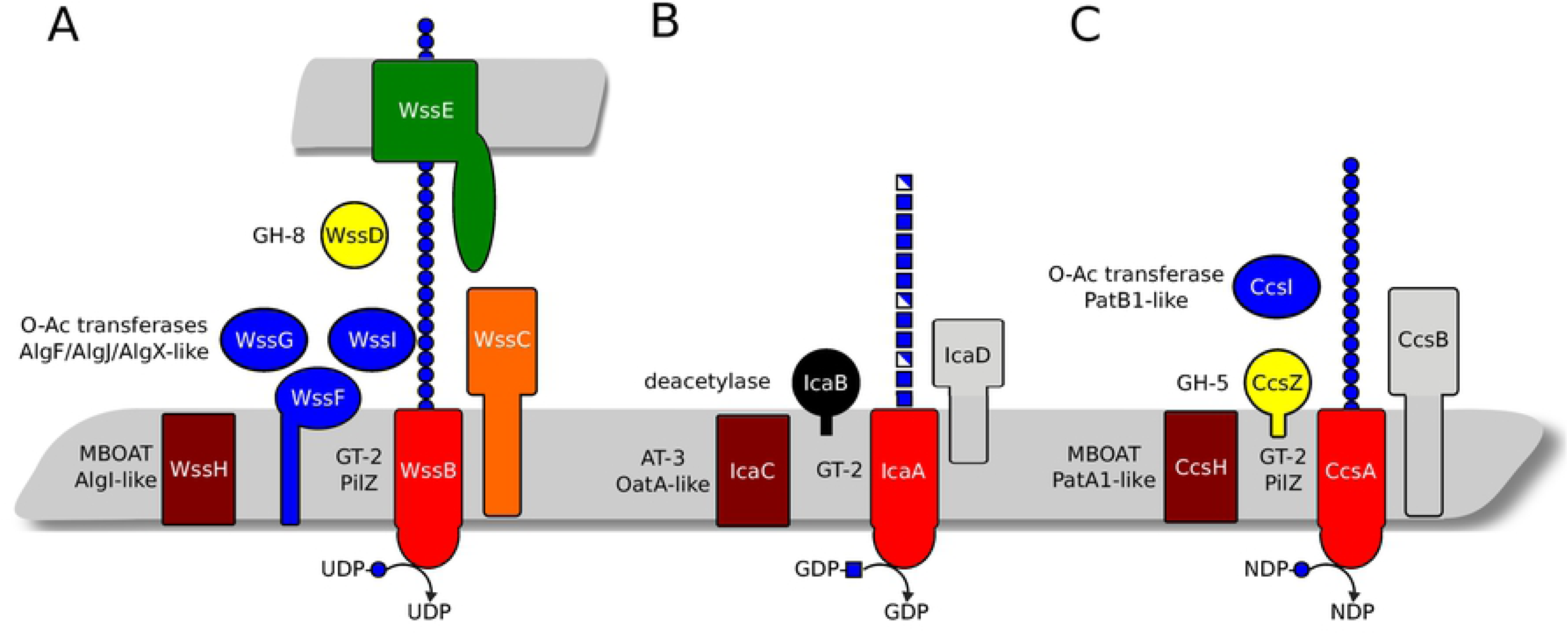
The *ccsABZHI gene cluster* shares structural similarity to both the Gram-negative O-acetylated cellulose synthase and the Gram-positive PNAG synthase. Proteins are coloured as in Fig. 1. (A) The model of the *P. fluorescens* SBW25 cellulose synthase based on Spiers *et al.* (2003; Ref.10), which contains putative O-acetyltransferases resembling those of the alginate biosynthesis pathway in *P. aeruginosa*. (B) The proposed model of PNAG synthesis by *S. aureus* and *S. epidermidis* based on Atkin *et al.* (2014; Ref. 26). Synthesis of PNAG depends on both IcaA and IcaD. O-succinylation is proposed to be carried out by IcaC, an AT-3 enzyme resembling OatA from *S. aureus*, and deacetylation is carried out by IcaB. (C) Our proposed model of the cellulose synthase from selected Clostridia. The synthases we identified contain the hallmark GT-2 and PilZ domains of Gram-negative cellulose synthases, in addition to CcsB, a membrane-bound extracellular protein of unknown function. O-acetylation of cellulose in this pathway is likely carried out by CcsH and CcsI, which resemble PatA1 and PatB1 from *B. cereus* and WssH and WssI from *P. fluorescens* SBW25, and cleavage is carried out by the GH-5 enzyme CcsZ reported here.

Although the identity of the exopolysaccharide in the *C. ventriculi* biofilm was originally identified as cellulose and not O-acetylated cellulose [8], it is worth noting that cells were treated with boiling sodium hydroxide during isolation of the exopolysaccharide from cultured biofilms, which may explain why O-acetylation evaded detection through decades of Clostridial biofilm research since ester-linked acetyl groups would not withstand this treatment. However, the production of O-acetylated cellulose by *C. ventriculi* and other Clostridia has yet to be experimentally validated, and the specific roles of CcsH and CcsI would also accordingly need to be demonstrated. We emphasize that in the absence of a complete structural and functional investigation of each of these individual proteins, our proposed model of the *ccsABZHI* synthase serves only as an entry point towards an understanding of cellulose biofilm formation by Gram-positive bacteria. To this end, we further examined the activity of CcsZ as a putative cellulose hydrolase.

### The overall structure of CcsZ

We engineered and synthesized the soluble domain of *ccsZ* from *C. difficile*, corresponding to residues 30-340 of the GenPept entry WP_077724661.1, into the pET21 expression vector to generate a recombinant His-tagged expression construct of CcsZ. Protein from this construct expressed well, yielding up to 75 mg/L of culture, and purified to near homogeneity following nickel affinity chromatography and anion exchange chromatography, making it an excellent candidate for structure determination. Our CcsZ crystals grew in space group *P* 2_1_ and diffracted X-rays well to 1.75 Å resolution with a single copy of CcsZ in the asymmetric unit. We solved the crystal structure of CcsZ using the molecular replacement technique based upon the model of apo-*Tm*Cel5A (PDB ID 3AMD) as a search template. Model building and subsequent refinement resulted in an improved *R*_work_ and *R*_free_ of 0.227 and 0.260, respectively. The model covers almost the entirety of the CcsZ construct, from residues 38 to 337 in the sequence, although residue Ser91 could not be reliably built into the model due to a lack of electron density. The complete data collection and refinement statistics are available in Table 1. The completed structure of CcsZ was deposited to the Protein Data Bank (PDB) under accession code 6UJE.

**Table 1:**
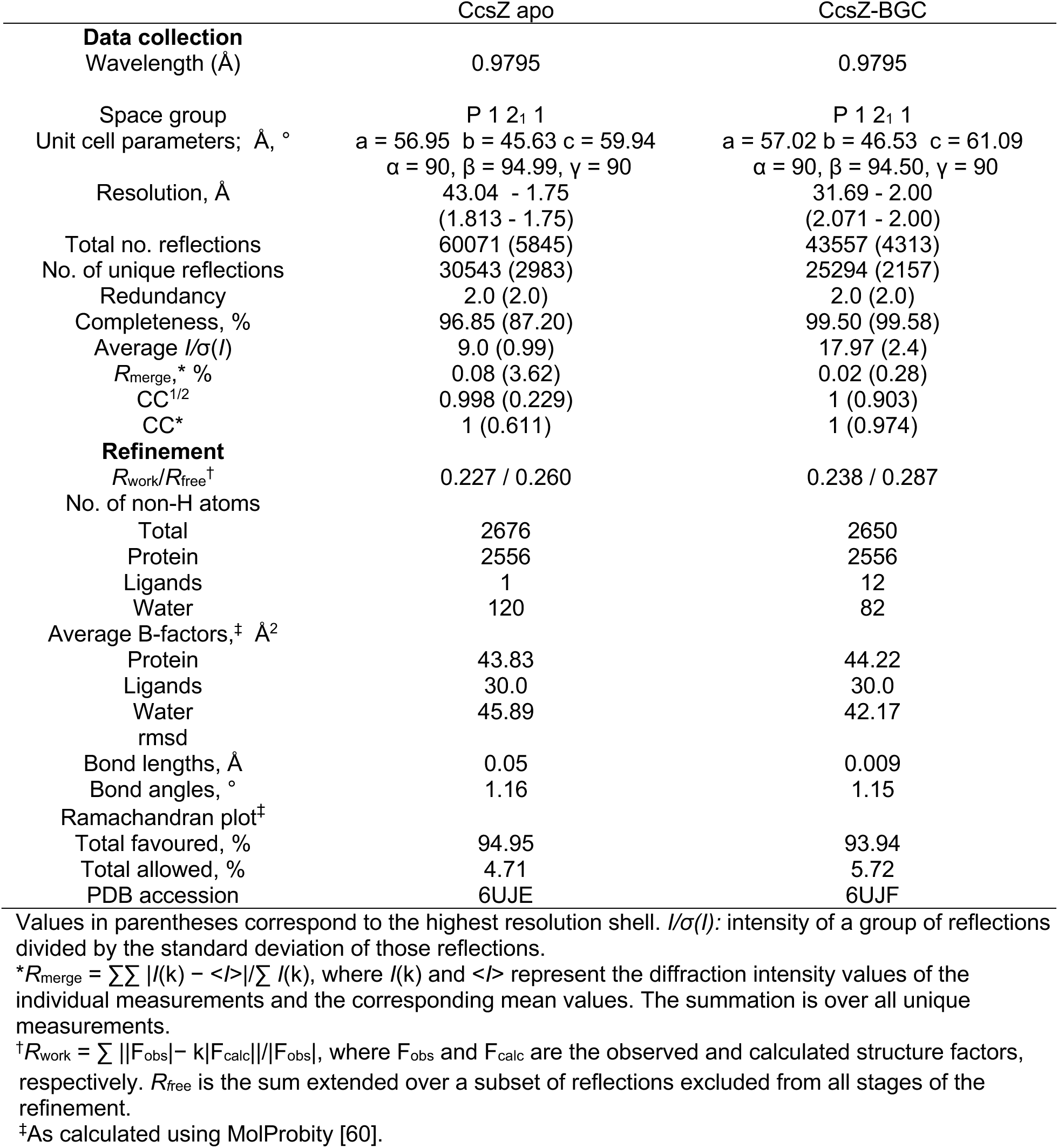
Diffraction data processing, refinement statistics, and model validation.

In agreement with typical GH-5 enzymes, CcsZ folds into an overall structure adopting a distorted TIM barrel fold, in which an (α/β)_8_ barrel is formed at the core of the fold by eight parallel β-strands (Fig. 3, A and B). This β-barrel motif is flanked by a series of eight partially distorted α-helices packed against the core β-strands, which are connected by extended loops along the C-terminal face of the β-barrel motif. The extended C-terminal loops shape a deep cleft that is typical of GH-5, where both the active site and the cleft for substrate accommodation are located. Interestingly, we observed this groove to be strongly negatively charged in CcsZ, which is not a typical feature of GH-5 (Fig. 3C). The GH-5 consensus catalytic acid/base and nucleophile residues are present in CcsZ as Glu-164 and Glu-281, respectively (Fig. 3A). Our structure of apo-CcsZ superimposed excellently on apo-*Tm*Cel5A, with an r.m.s.d. of 0.677 Å across 252 equivalent Cα atoms, as expected given the reasonably high sequence identity (43%) between these proteins and the highly conserved GH-5 fold.

**Figure 3.**
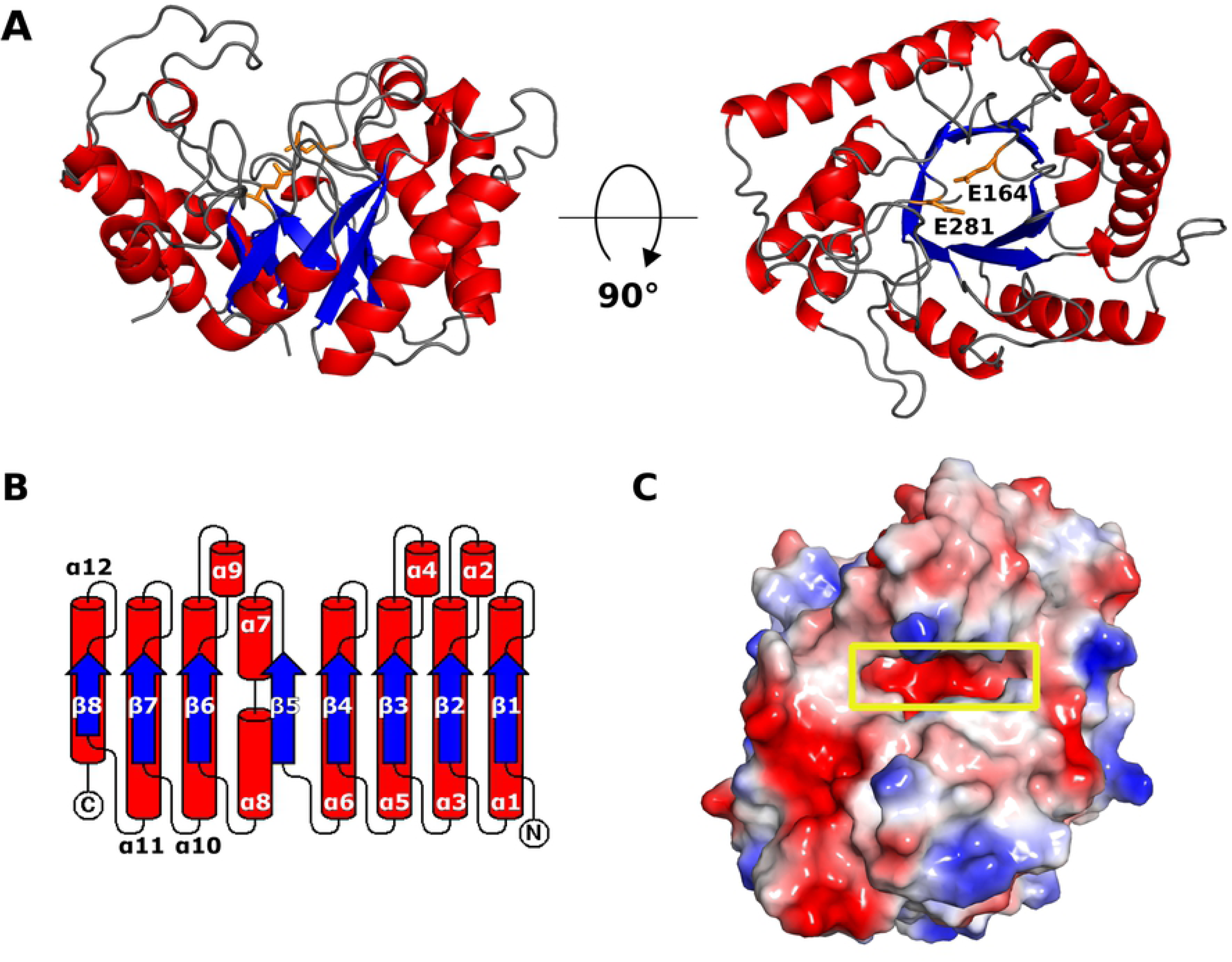
The overall structure of CcsZ. (A) The (α/β)_8_ fold of CcsZ is shown as a cartoon representation from a front (left) and top (right) view. The catalytic resides Glu-164 and Glu-281 are shown in orange. (B) Topology cartoon of the CcsZ fold, coloured as in panel A. (C) The substrate-binding cleft of CcsZ (yellow box) has an overall strong negative charge, an unusual feature for GH-5 enzymes.

### Comparison to known GH-5 structures

Carbohydrate-Active Enzyme family 5 (CAZy GH-5) is an exceptionally large family containing >15,000 known and predicted enzymes. In keeping with the size of GH-5, the substrate specificity of this family is widely variable and encompasses numerous carbohydrate structures, with some members demonstrating strict specificity for single glycan structures, while other members have demonstrated promiscuity with activity on many distinct glycans [49]. The diversity of substrate preference in the GH-5 family is also matched by the wide variety of chemical properties among GH-5 enzymes, such as temperature or pH optima [49–53]. In efforts to delineate the diversity of this large family of glycosyl hydrolases, phylogenetic analyses were performed and numerous sequence-based phylogenetic subfamilies with characteristic substrate preferences were created [49]. Under this classification, Aspeborg *et al.* placed CcsZ into GH-5 subfamily 25 (GH-5_25), a small subfamily of enzymes belonging primarily to thermophilic bacteria, although two other GH-5_25 enzymes from *C. difficile* were also classified in GH-5_25 [49]. It was not surprising that based upon the molecular phylogenetics of GH-5_25, the *Tm*Cel5A enzyme we used as the search model for molecular replacement of CcsZ is also among the most closely related to CcsZ of all GH-5 sequences [49].

GH-5_25 is reported as a polyspecific subfamily of GH-5 that possesses multiple activities [49]. Notably, *Tm*Cel5A was reported to exhibit activity on both mannan and glucan polymers with β-(1,4) linkage, including linear and branched polysaccharides [54]. Supporting this finding, the crystal structure of apo-*Tm*Cel5A was also reported alongside structures of *Tm*Cel5A ligand complexes, including *Tm*Cel5A bound to cellotetraose, mannotriose, glucose and cellobiose. We compared the structure of *Tm*Cel5A in complex with cellotetraose (PDB ID 3AZT) to CcsZ in an effort to understand how their substrate accommodation might differ. The *Tm*Cel5A-cellotetraose complex also superimposed well over CcsZ, with an r.m.s.d. of 0.620 Å over 242 equivalent Cα atoms. A comparison of the substrate-binding cleft, informed by the positioning of the cellotetraose ligand of *Tm*Cel5A, demonstrated a clear difference between the subsite architectures of the two enzymes (Fig. 4). The residues that contribute to the −1 and −2 subsites of *Tm*Cel5A appear to be partially conserved in CcsZ, present as H123, H124, N163, Y225, and N48. However, in *Tm*Cel5A, W210 shapes the face of the −2 subsite and stacks with glucopyranose rings, but in CcsZ, the equivalent Y237 occupies what would serve as the −3 subsite in *Tm*Cel5A; thereby resulting in a substrate binding cleft in CcsZ that terminates at the equivalent −2 subsite.

**Figure 4.**
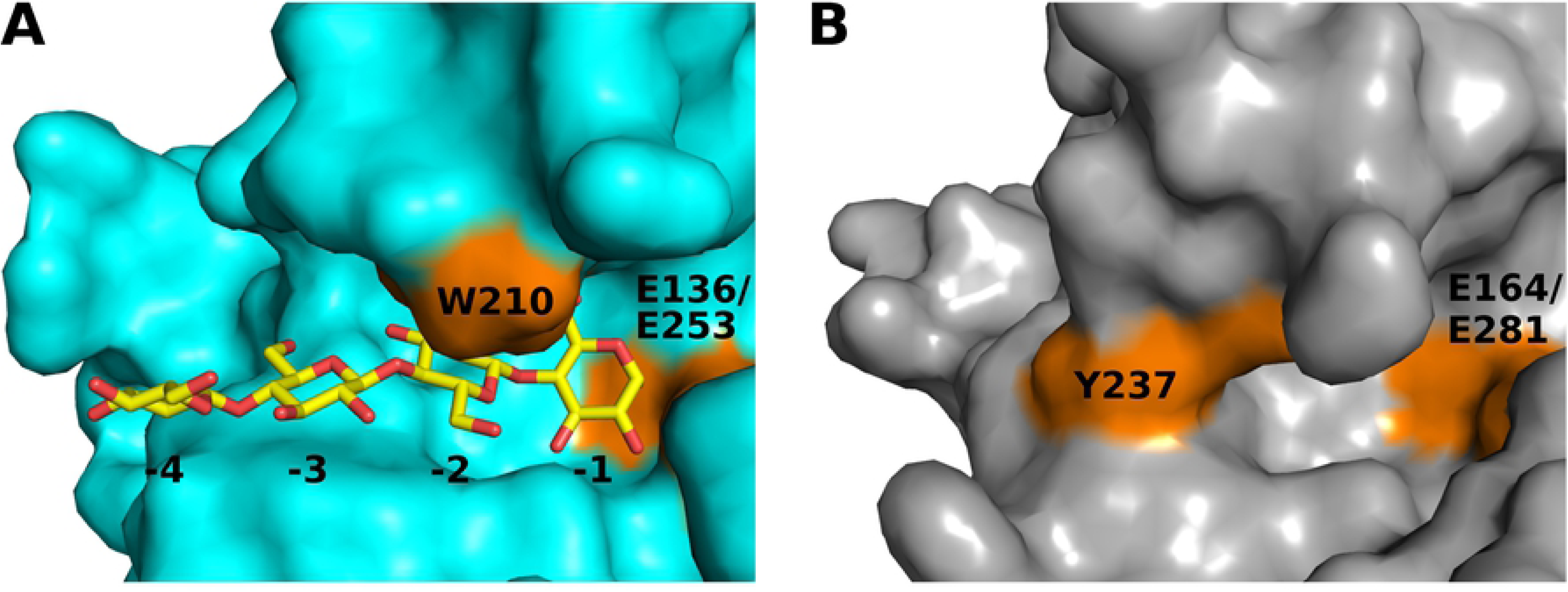
Substrate accommodation by CcsZ and *Tm*Cel5A. (A) The cellotetraose-bound structure of *Tm*Cel5A. The cellotetraose molecule is bound at subsites adjacent to the active site residues Glu-136 and Glu-253, guided by ring-stacking with Trp-210. (B) The equivalent substrate-binding site of CcsZ. The active site residues are present as Glu-164 and Glu-281. The equivalent to the −2 subsite, shaped by W210 in *Tm*Cel5A, is obstructed by the orientation of Tyr-237 in CcsZ, preventing substrate accommodation in the same fashion.

*Tm*Cel5A was also reported to be highly thermostable [53], a feature rationalized by a larger fraction of buried atoms, a smaller accessible surface area, and the presence of shorter unstructured loops as compared to the structure of a mesophilic GH-5 cellulase from *Clostridium cellulolyticum* (*Cc*Cel5A; PDB id 1EDG) [50]. However, for the purpose of comparison, it is worth noting that *Cc*Cel5A does not belong to GH-5_25 but instead to GH-5_4, which includes *endo*-β-(1,4) glucanases specific for xyloglucan, as well as licheninases and xylanses. To assess if CcsZ possessed similar features attributed to the thermal stability of *Tm*Cel5A, we uploaded our structure to the PDBePISA server. This analysis revealed both a solvent-accessible surface area for CcsZ of 13497 Å^2^ and a proportion of buried atoms of 0.50 that was most similar to *Tm*Cel5A (13720 Å^2^ and 0.51) rather than *Cc*Cel5A (15,410 Å^2^ and 0.46). Furthermore, CcsZ possessed unstructured loops more similar to *Tm*Cel5A (r.m.s.d. 0.677 Å across 252 Cα atoms) than *Cc*Cel5A (r.m.s.d. 1.053 Å across 172 Cα atoms) based on superimposition of the structures.

After comparison of our structure of CcsZ to other related GH-5 structures, we found the relatively high sequence identity and structural similarity between CcsZ and *Tm*Cel5A rather than *Cc*Cel5A (i.e. 41% vs 18% id) surprising, based on our expectation that CcsZ would be highly specific for cellulose or possibly O-acetylated cellulose, in contrast to the broad specificity of *Tm*Cel5A for varied mannan and glucan structures. We also did not anticipate CcsZ to possess features associated with extended thermal stability given that *C. difficile* is a mesophilic bacterium and that the biofilm phenotype has been reported at temperatures of 25-37°C [8, 32]. Accordingly, we set out to biochemically characterize CcsZ to further explore these properties.

### The product-bound structure of CcsZ

Next, in order to experimentally resolve the subsite architecture and mechanism of substrate accommodation by CcsZ, we attempted to solve the structure of CcsZ in complex with cello-oligosaccharides. We were able grow crystals in a distinct but chemically similar condition to our apo-CcsZ crystals and successfully introduced cellotriose following crystal growth but prior to crystal harvesting. These crystals diffracted X-rays to 1.65 Å resolution and also grew in the space group *P* 2_1_ containing a single polypeptide in the asymmetric unit (Table 1). We solved the structure of our ligand complex with molecular replacement using the structure of the apo-form as the search model. The ligand-bound structure was in complete agreement with the apo-form, and although the experimental resolution was higher for the ligand complex crystals than the apo-form, low completeness was observed for the high-resolution data and so the ligand complex structure was refined only against data with a maximum resolution of 2.0 Å. Refinement of the ligand complex structure resulted in an *R*_work_ and *R*_free_ of 0.238 and 0.287, respectively (Table 1). The final model of the CcsZ-ligand complex also covered almost the entirety of the protein, from residues 39-340, and included the missing S91 from the apo-form.

Following refinement of the complex structure, we observed a Fourier electron density peak, clearly interpretable in both the Fourier 2 *F_o_ - F_c_* electron density map at 1.0 σ contour level and the *F_o_ - F_c_* map at 3.0 σ contour level. This density corresponded well to an apparent glucopyranose ring near the active site. Subsequent ligand fitting placed glucose into this density with a promising real-space correlation coefficient (RSCC) of 0.714 in an unambiguous orientation, with placement aided by the strong electron density observed for the oxygen atom of the C6 hydroxyl. We then calculated a Polder omit map for the glucose ligand, resulting in an improved electron density that allowed refinement of the glucose ligand to a final RSCC of 0.861 (Fig. 5C). The bound glucose makes polar contacts with the side-chain N atom of N48 via the C2 hydroxyl, the N atom of the H123 side chain via the C6 hydoxyl, and the N atom of the W314 side chain via the C6 hydroxyl, with approximate distances of 3 Å in all cases (Fig 5B). The final refined structure of the CcsZ-glucose complex was deposited to the PDB under accession code 6UJF.

**Figure 5.**
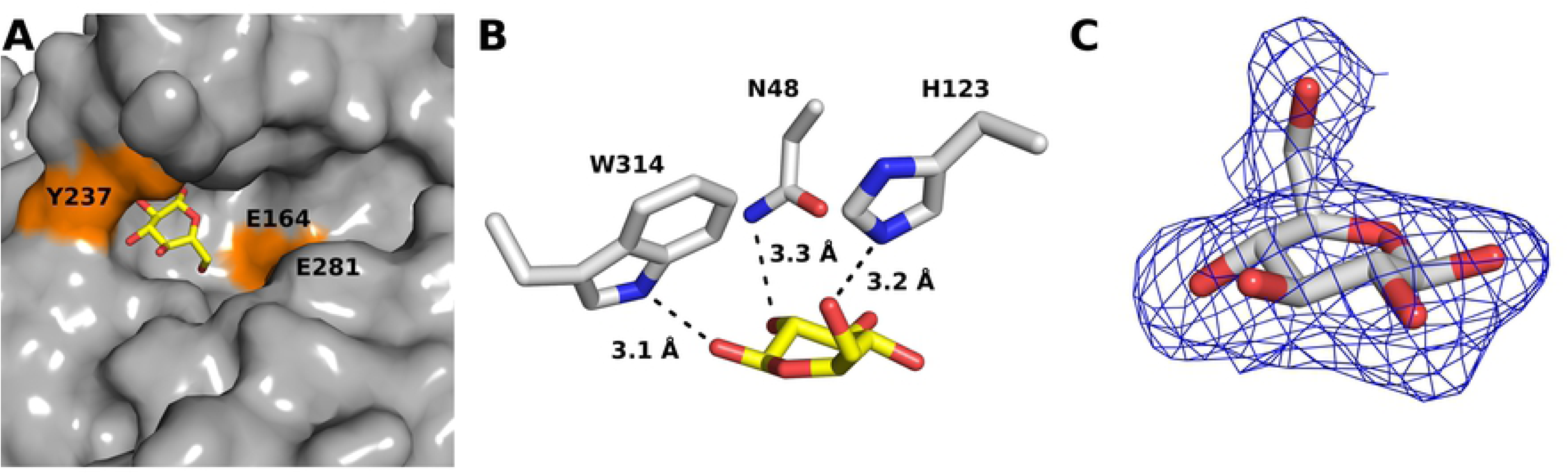
The product-bound structure of CcsZ. (A) A glucose molecule is bound at the apparent +1 subsite adjacent the catalytic architecture and confined by the orientation of Y237. (B) The bound glucose molecule makes polar contacts with W314 via the C1 hydroxyl and N48 via the C2 hydroxyl, and H123 via the C6 hydroxyl. (C) The polder omit map electron density (blue mesh) for the glucose ligand contoured at 3 σ.

The observation of a single glucose residue bound to CcsZ when a trisaccharide was soaked into the crystals suggested to us that the bound ligand represented an enzymatic product rather than a bound substrate or glucosyl intermediate, which was not surprising given the observed activity of our CcsZ construct specifically on glucan substrates (noted below). Accordingly, this glucose molecule was bound at the equivalent of the −2 subsite in *Tm*Cel5A, but appears spatially confined by the side chain of Y237 and the loop presenting W314, both of which shifted only trivially between our apo- and glucose-bound structures with an all-atom r.m.s.d. of 0.206 Å across 2232 equivalent atoms (Fig. 5A). Comparatively, the Gram-negative cellulose hydrolase BcsZ (a GH-8 enzyme), which was resolved both in apo-form and as a complex with cellopentaose, displays a substrate-binding architecture whereby the +1 and +2 subsites are angled at approximately 60° from the −1 to −4 subsites at the nonreducing side of the catalytic center. Thus, structural plasticity of the substrate-binding groove and induced-fit distortion of longer saccharide substrates is certainly possible in these enzymes and is also conceivable in CcsZ, given the structure we observed.

### CcsZ is an *endo*-β-glucanase

To test if CcsZ was in fact capable of hydrolytic activity on cellulose, we assessed its activity on the soluble substrate analogue carboyxmethylcellulose (CMC) first using the CMC-agar overlay method reported by Mazur and Zimmer for BcsZ [27]. Following staining with Congo Red, a zone of clearing was observed suggesting CcsZ was capable of cleaving the glucosidic bonds of CMC after 1 h incubation (Fig 6A). We also performed this same agar overlay experiment using equal concentrations (0.8% w/v each) of CMC, hydroxyethyl cellulose (HEC) and beechwood xylan with a 24 h incubation. As expected, we observed that CcsZ was active on CMC and HEC but was only capable of partially cleaving xylan (Fig. S1).

**Figure 6.**
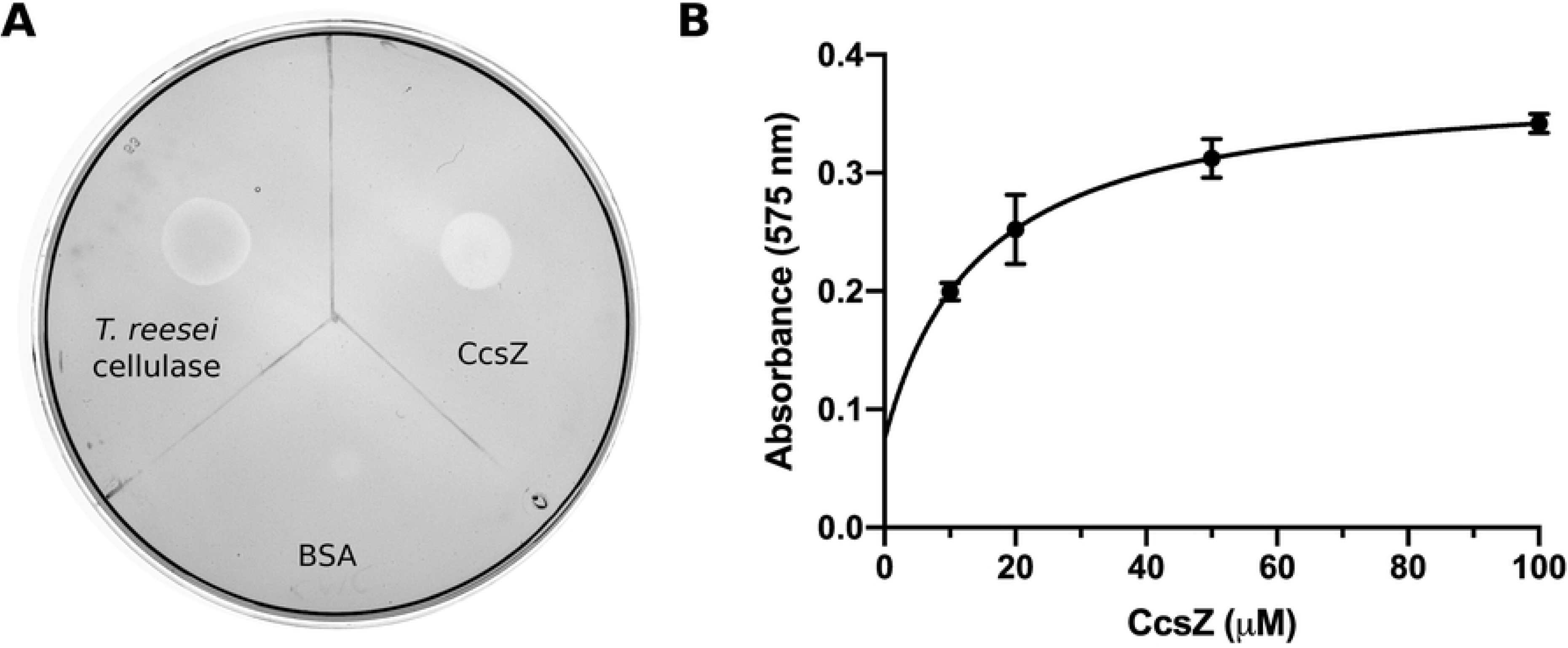
CcsZ is an *endo*-β-glucanase. (A) CMC hydrolysis by CcsZ and the *T. reesei* cellulase cocktail positive control is evident by the zone of clearing in the Congo Red stained CMC-agar overlay. (B) CMC hydrolysis by CcsZ. The reducing sugar concentration of a CMC solution (16.6 g/L) was increased by treatment with CcsZ over 4 h at 37°C in a CcsZ concentration-dependent manner.

Subsequently, we sought to measure CcsZ activity on CMC quantitatively using the dinitrosalicylic acid (DNS) reducing sugar assay, where enzyme activity was calculated from our raw data using a standard curve of glucose prepared using the DNS method. We observed an increase in reducing sugar concentration following incubation with CcsZ in a concentration-dependent manner, further indicating CcsZ is capable of using CMC as a substrate (Fig. 6B). We also tested CcsZ with CMC under a range of pH buffer conditions to measure pH stability. We found CcsZ to have a pH optimum of approximately 4.5, which is typical of other characterized GH-5 enzymes (Fig. 7A) [50–52]. Although CcsZ was most active under acidic conditions, we found CcsZ was tolerant of a wide variety of pH and buffer conditions and still demonstrated detectable hydrolase activity on CMC across all pH values we tested (pH 4-10). CcsZ activity was reduced two-fold between pH 4.5 and 7.5, with a particular loss in activity under alkaline conditions, showing a six-fold reduction in activity at pH 9.5.

**Figure 7.**
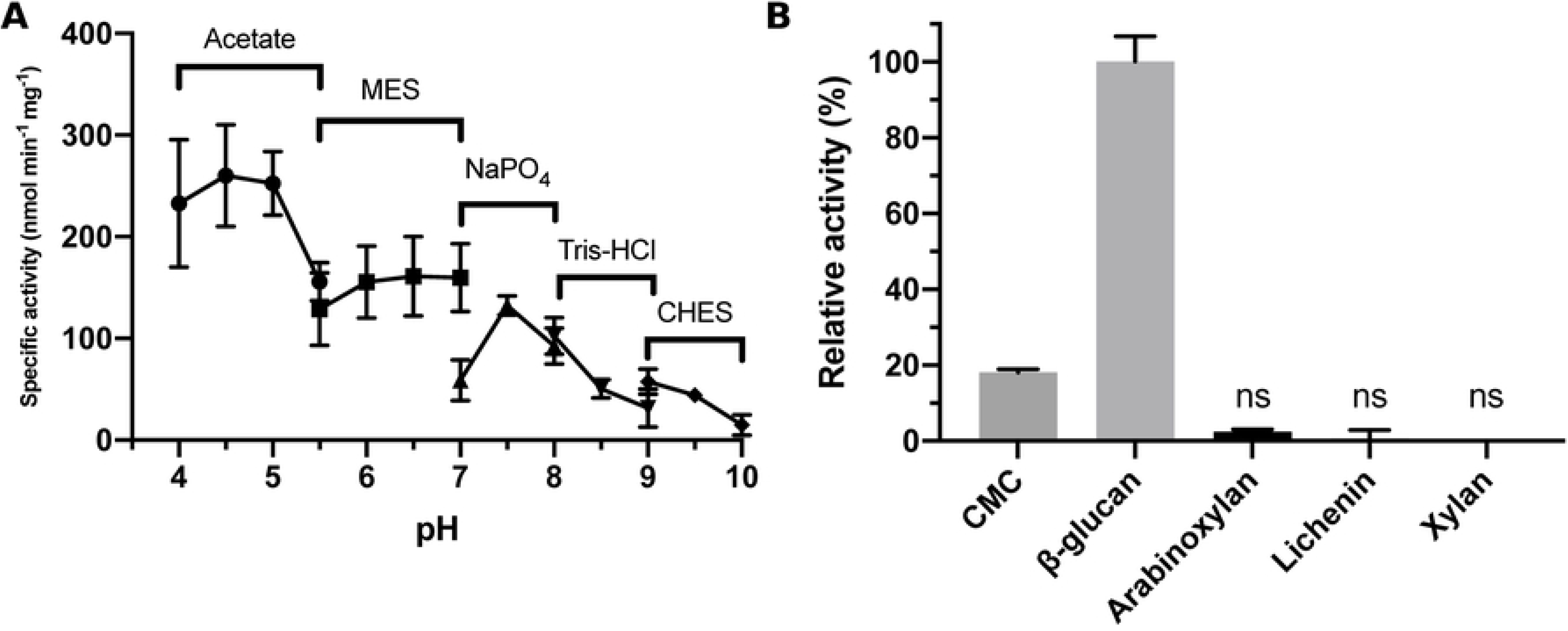
pH and substrate specificity of CcsZ. (A) The pH profile of CcsZ displays a clear preference for acidic conditions with a pH optimum of 4.5. Activity was calculated using the DNS method with solubilized CMC substrate. Buffer solutions used (50 mM) are listed above data points. (B) Substrate utilization profile of CcsZ using common GH-5 substrates. CcsZ exhibited five-fold greater activity on mixed-linkage β-glucan as compared to CMC when assayed using the DNS method. CcsZ activity on arabinoxylan, lichenin, and xylan was not significantly different from an enzyme-free control under our assay conditions.

To assess substrate specificity, we also tested the common GH-5 polysaccharide substrates arabinoxylan, xylan, lichenin and β-glucan using the same DNS reducing sugar assay. We did not observe a significant increase in reducing sugar concentration following prolonged CcsZ incubation (*i.e.* 24 h) with arabinoxylan, xylan, or lichenin under our conditions (Fig. 7B), consistent with the expected activity of CcsZ on cellulose exopolysaccharides and its classification in GH-5_25. However, we observed CcsZ exhibited 5-fold greater activity on β-glucan, a polymer with mixed β-(1,3) and β-(1,4) linkages. In agreement with our structural data, this observation suggested that carboxymethylcellulose was actually a poor CcsZ substrate owing to the stearic constraints of the carboxymethyl groups. Instead, activity on a β-glucan is likely more representative of the minimal, unbranched structure of cellulose exopolysaccharides, and others have also reported that β-glucan is a superior substrate mimetic to CMC for GH-5 strict *endo-*β glucanases [51].

*Exo*-acting activities are rare among GH-5 enzymes, with the majority of characterized GH-5 enzymes characterized as *endo-*acting, although it is worth noting that this is a challenging biochemical distinction to establish experimentally, and that these activities may not be mutually exclusive. In the case of CcsZ, an unusual substrate-accommodation cleft in addition to our structure of CcsZ bound to a glucose monosaccharide suggested that CcsZ may exhibit strict *exo*-glucanase activity, which warranted further investigation. To assess the regioselectivity of CcsZ, we incubated the enzyme with the mock substrate cellopentaose (G_5_) and analyzed the enzymatic products by liquid chromatography-mass spectrometry (LC-MS; Fig. 8 and S2, Table 2). Our enzyme-free control contained only the m/z species for the intact G_5_ starting material, indicating the substrate was not solvent-labile. In the mass spectra of the CcsZ enzymatic products, we observed predominantly m/z species that corresponded to the cellobiose disaccharide (G_2_) and cellotriose trisaccharide (G_3_) species. In addition, we also observed unreacted and intact starting material (G_5_) among the CcsZ products, as well as m/z species that corresponded to the glucose monosaccharide (G_1_) and cellotetraose oligosaccharide (G_4_). Although not truly quantitative, we noticed the relative abundance of these m/z species was markedly lower than that of the di- and trisaccharide species, and that their intensity on the liquid chromatograph matched this low ionization potential under constant conditions. This data suggested the preferred regioselectivity of CcsZ as an *endo*-glucanase and also accounted for the presence of small quantities of glucose, likely an enzymatic side product, which we were able to structurally resolve bound to CcsZ.

**Figure 8.**
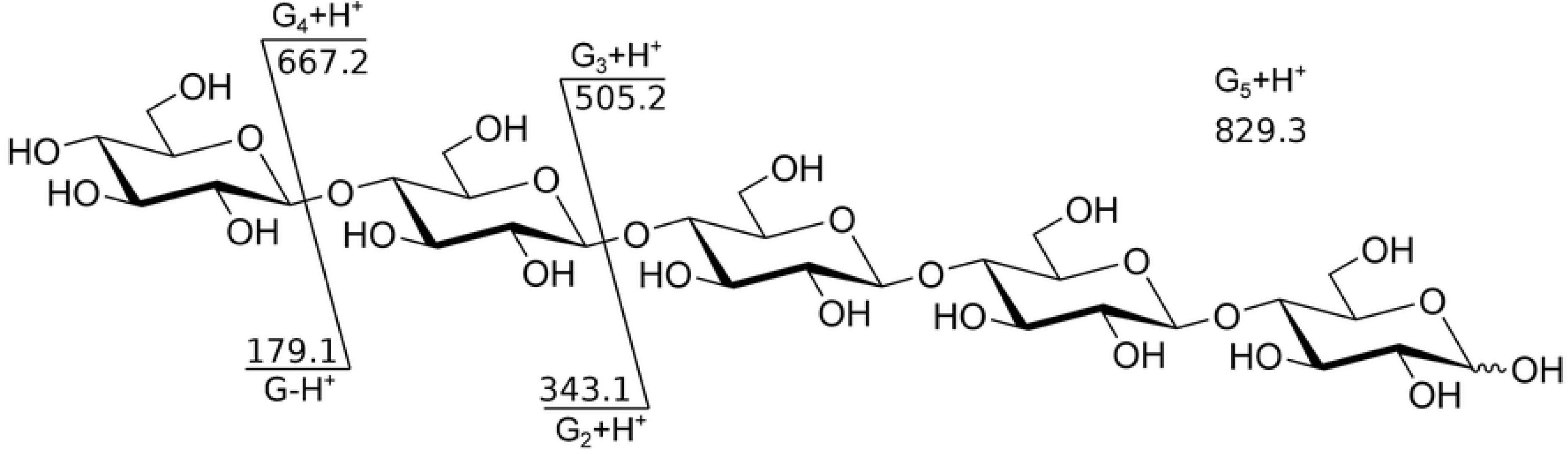
The observed m/z species and corresponding structure of the CcsZ enzymatic products. Cleavage of cellopentaose (G_5_) by CcsZ was observed to occur in an *endo*-acting fashion, resulting in products G_2_ and G_3_. Additionally, enzymatic products G_1_ and G_4_ were observed in lower relative abundance, corresponding to *exo*-acting hydrolytic activity at terminal saccharides.

**Table 2:** CcsZ enzymatic products from G_5_ substrate detected by LC-MS.

| Species | Formula | RT | Observed Mass (amu) | Rel. Abundance | Calculated Mass (amu) | Difference(ppm) |
| --- | --- | --- | --- | --- | --- | --- |
| G<sub>1</sub> | C<sub>6</sub>H<sub>12</sub>O<sub>6</sub> | 0.803 | 180.0645 | 1653 | 180.0634 | 6.44 |
| G<sub>2</sub> | C<sub>12</sub>H<sub>22</sub>O<sub>11</sub> | 1.045 | 342.1166 | 3561 | 342.1162 | 1.15 |
| G<sub>3</sub> | C<sub>18</sub>H<sub>32</sub>O<sub>16</sub> | 1.045 | 504.1691 | 29206 | 504.169 | 0.13 |
| G<sub>4</sub> | C<sub>24</sub>H<sub>42</sub>O<sub>21</sub> | 1.07 | 666.2211 | 555 | 666.2219 | -1.15 |
| G<sub>5</sub> | C<sub>30</sub>H<sub>52</sub>O<sub>26</sub> | 1.021 | 828.2742 | 714 | 828.2747 | -0.56 |

### Conclusions

The class Clostridia contains numerous pathogens of global health importance. *Clostridioides difficile* is one of many such pathogens that poses a global threat to public health. *C. difficile* infection poses significant morbidity and mortality to all populations worldwide and among individuals beyond the groups traditionally recognized at-risk (*e.g.* the elderly, those under hospital care, or those under antimicrobial therapy) [55–57]. The burden of *C. difficile* diarrheal disease is also not limited to developing nations: in the U.S., *C. difficile* caused an estimated 453,000 infections resulting in 29,300 deaths in 2012 alone [56]. Although healthcare-associated costs are challenging to estimate, this burden of *C. difficile* infection results in as estimated cost of approximately US $6 billion dollars per year in the United States alone. This data is notwithstanding the other important human pathogens belonging to class Clostridia. Biofilms have demonstrated roles in bacterial persistence and virulence [12,13,58,59] and have been anecdotally reported in Clostridia to be composed of cellulose exopolysaccharides [7, 8]. Additionally, the loci encoding the molecular machinery for Pel polysaccharide has been recently identified in members of Clostridia [33]. Here, we demonstrate the presence of the *ccsABZHI* locus that bears sequence similarities to known Gram-negative cellulose exopolysaccharide biosynthesis systems. Our subsequent analysis of the cognate glycosyl hydrolase, CcsZ, reveals the Clostridial cellulose synthase contains a functionally analogous but structurally distinct glycosyl hydrolase from the known Gram-negative cognate glycosyl hydrolase BcsZ. The findings we present here represent an entry point to an understanding of the molecular mechanisms governing cellulose exopolysaccharide production in Gram-positive bacteria. More specifically, this work verifys the ability of Clostridia to produce cellulose-containing biofilms and the underlying results form the basis for future work to improve the control of problematic infections caused by these bacteria.

## Acknowledgements

We thank Dr. Dyanne Brewer and Dr. Armen Charcoglycan at the Mass Spectrometry facility of the Advanced Analysis Center, University of Guelph for their technical assistance and expertise with mass spectrometry experiments. JTW is supported by the Natural Sciences and Engineering Research Council of Canada (NSERC) in the form of a grant (#418310). AA was previously supported by an Ontario Graduate Scholarship via Wilfrid Laurier University. Research described in this paper was performed using beamline 08ID-1 at the Canadian Light Source, which is supported by the Canada Foundation for Innovation, Natural Sciences and Engineering Research Council of Canada, the University of Saskatchewan, the Government of Saskatchewan, Western Economic Diversification Canada, the National Research Council Canada, and the Canadian Institutes of Health Research.

## Supporting Information Figure legends

**Figure S1. CcsZ activity assessed using a semisolid agar-carbohydrate assay containing 0.8% (w/v) each CMC (A), HEC (B), and beechwood xylan (C).** CcsZ was capable of complete CMC hydrolysis, resulting in a total loss of Congo Red staining where spotted on the agar, but only partial degradation of HEC and xylan were observed.

**Figure S2. Summary of CcsZ enzymatic products detected by LC-ESI-MS.** All possible cleavage products of cellopentaose (G_5_), G_1_ (A), G_2_ (B), G_3_ (C), G_4_ (D) were detected, in addition to unmodified starting product G_5_ (E).

